# Polycomb repressive complex 2 regulates sexual development in *Neurospora crassa*

**DOI:** 10.1101/2025.05.15.654408

**Authors:** Abigail M. Deaven, Abigail J. Ameri-Solanky, Zachary A. Lewis

## Abstract

In most branches of the fungal kingdom, sexual development marks a dramatic shift from simple multicellular hyphae to complex fruiting bodies composed of multiple tissues and cell types. It is essential to tightly regulate development to prevent unnecessary energy investment into building these structures. Polycomb Repressive Complex 2 (PRC2) is a highly conserved regulator of development in multicellular organisms. In plants, animals, and some fungi, PRC2 tri-methylates histone H3 lysine 27 in promoters and gene bodies to repress gene expression. In *Neurospora crassa*, H3K27me3-associated genes are poorly conserved and stably repressed in standard lab conditions. Through analysis of publicly available RNA-seq experiments, we found that PRC2-methylated genes are broadly and uniquely activated during sexual development. PRC2-methylated genes comprise a distinct subset of developmentally induced genes (DIGs) characterized by an exceptionally high degree of cell-type specificity. Loss of PRC2 activity results in the precocious formation of perithecia-like structures (dubbed false perithecia), even in the absence of a compatible mating-type partner. These structures show a unique gene expression profile with activation of both PRC2-methylated and unmethylated DIGs, showing evidence of a transcriptional reprogramming event. Together, these data suggest that PRC2 is part of a developmental checkpoint that restricts fruiting body development to the appropriate conditions.

## Importance

Development of multicellular eukaryotes involves transcriptional reprogramming to drive cell fate transitions. This study identified PRC2 as a critical regulator of cell fate in the model filamentous fungus *Neurospora crassa*, where it silences a subset of sexual development genes. Loss of regulation by PRC2 triggers a major reprogramming event in which genes specifying sexual tissues cannot be repressed, causing a homeotic transition. These results provide novel insights into the role of PRC2-mediated regulation in the fungal kingdom and uncover a critical checkpoint regulating complex multicellular development.

## Observation

Across eukaryotes, key developmental transitions are regulated by a complex network of transcriptomic, proteomic, and epigenetic regulatory pathways. A central player in this regulation is Polycomb Repressive Complex 2 (PRC2), a conserved chromatin modifier that catalyzes tri-methylation of histone H3 lysine 27 (H3K27me3) to stably repress genes (1). In plants, insects, and animals, defects in H3K27me3-dependent repression are linked to homeotic transformations, underscoring PRC2’s essential role in organismal development (2–5). Despite its well-established role in development of higher eukaryotes, the biological significance of PRC2-mediated repression is poorly understood in fungi.

PRC2 and H3K27me3 are absent from the widely used model yeasts, *Saccharomyces cerevisiae* and *Schizosaccharomyces pombe*, but genes encoding PRC2 components are conserved in many fungal lineages (6). The first functional studies of a fungal PRC2 complex were carried out in *Neurospora crassa*, where approximately 7% of genes are enriched for H3K27me3 (7). These genes are tightly repressed in standard growth conditions, and genetic studies have shown that repression depends on the histone variant H2A.Z (8), the chromatin remodeler Imitation Switch (ISW) (9, 10), the H3K27me3 reader EPR-1 (11), the RPD3L complex (12), the PRC2 Accessory Subunit (PAS) (13), and components of the constitutive heterochromatin pathway (14–16). While mechanistic insights into H3K27me3-dependent repression have been gained, the biological functions of PRC2 and PRC2-repressed genes in *N. crassa* have remained mysterious for more than a decade. H3K27me3-enriched domains are overrepresented for poorly conserved genes that are genus- or species-specific and lack conserved functional domains (7). Moreover, PRC2-deficient strains have modest phenotypes. Strains lacking H3K27me3 show normal vegetative growth rates and normal development of asexual spores (conidia), sexual fruiting bodies (perithecia), and ascospores (7). While the functions of H3K27me3-enriched genes in *N. crassa* are unknown, work in other fungal systems has linked H3K27me3 and PRC2 components to diverse processes including plant pathogenesis, symbiosis, and secondary metabolism (17–24). To understand the diversity of PRC2-dependent gene regulation within the fungal kingdom, we sought to investigate PRC2 function in *N. crassa*.

PRC2-methylated genes are not expressed in standard lab conditions. We hypothesized that PRC2-methylated genes can be expressed under specific conditions and that expression conditions might provide insights into the biological functions of PRC2 and PRC2-repressed genes. To test this, we obtained 428 publicly available RNA-seq experiments performed with wild-type *N. crassa* grown in a variety of nutritional conditions and at different stages of asexual or sexual development (Figure 1a; Table S1; Table S2). We then compared the relative expression of known H3K27me3-enriched genes in each condition. For each experimental condition, we calculated a “gene induction score” for PRC2 methylated genes by multiplying the average expression level of all PRC2-methylated genes by the percent of PRC2-methylated genes induced (Figure 1b). This analysis revealed a striking upregulation of PRC2-methylated genes under two experimental conditions: when Avicel was used as the sole carbon source, and during multiple stages of sexual (perithecial) development (Figure 1b; Figure 1c). We next plotted expression of individual PRC2-methylated genes in each experimental condition (Figure 1d). Coordinated induction of most PRC2-methylated genes occurred exclusively during perithecial development, though subsets of PRC2 target genes were weakly induced in other conditions such as treatment with cell wall-inhibiting drugs and growth on Avicel. Induction of PRC2-methylated genes during perithecial development was confirmed in a second RNA-seq dataset (Figure S1).

**Figure 1.**
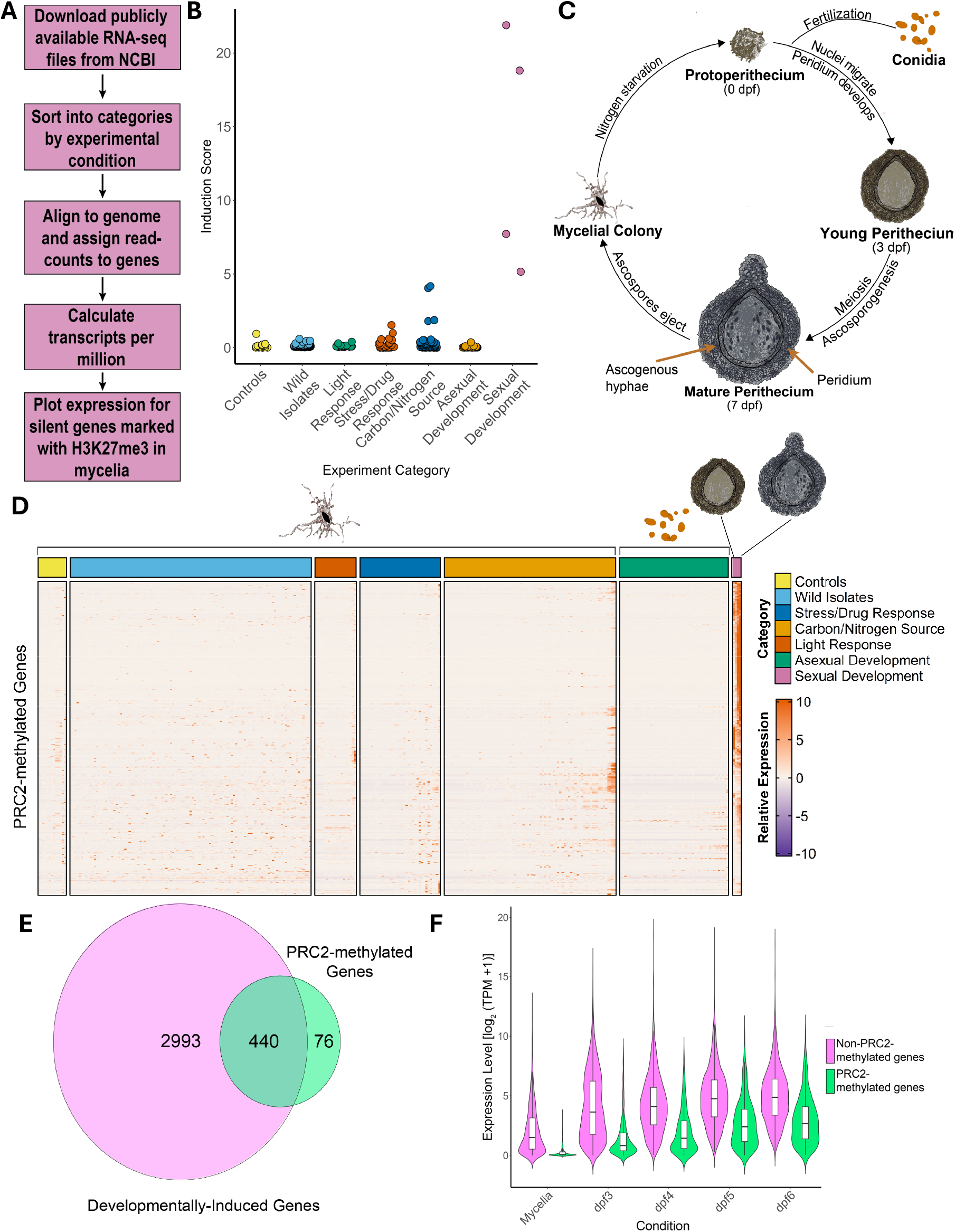
PRC2 methylated genes are broadly induced during sexual development. **(A)** The bioinformatics workflow used to examine expression of H3K27me3-enriched genes is shown. (B) The jitterplot shows the induction score of H3K27me3-enriched genes (y-axis) in each RNA-seq experiment (points). “Induction score” is the product of the mean expression of all H3K27me3-enriched genes in each experiment (TPM+1) and the percentage of H3K27me3-enriched genes with a z-score greater than 0.5. (C) The schematic illustrates *N. crassa* sexual development. (D) The heatmap depicts relative expression of individual H3K27me3-enriched genes (rows; n = 516) in publicly available RNA-seq experiments (columns). (E) The Venn diagram shows overlap between genes induced during sexual development (l_2_fc > 2, FDR < 0.05; n = 2,993) and genes enriched for H3K27me3 (n = 516. (E) The violin plot depicts the expression level (log_2_[TPM+1]) of DIGs that are enriched for H3K27me3 (green; n = 474) and DIGs that are not associated with H3K27me3 (magenta; n = 2,791) in mycelia or in developing perithecia isolated at 3, 4, 5, or 6-days post-fertilization (dpf).

To further explore the relationship between perithecial development and PRC2-methylated genes, we performed differential expression analysis to identify all genes that are induced in developing perithecia. We identified a total of 2,993 developmentally induced genes (DIGs), representing ∼1/3 of the *N. crassa* genome and 85% (440/516) of H3K27me3-enriched genes (Figure 1d; Table S3). We next compared expression of PRC2-methylated DIGs and unmethylated DIGs across developmental stages (Figure 1e). H3K27me3-marked DIGs are strongly repressed in mycelial tissue, while DIGs that lack H3K27me3 are expressed in both mycelia and perithecia. We conclude that PRC2 regulates perithecial-specific genes in *N. crassa*.

Because H3K27me3 directs formation of repressed chromatin, we hypothesized that PRC2 represses fruiting body development. Consistent with this idea, a previous study showed that PRC2-deficient mutants produce false perithecia, which were defined as melanized structures larger than 85 µm that form in the absence of fertilization (11). To determine if the false perithecia produced in PRC2-deficient strains represent aberrant perithecial-like structures, we first characterized the numbers and sizes of melanized structures produced by the unfertilized Δ*set-7* mutant, which lacks the catalytic subunit of PRC2. We measured the diameter of melanized structures that formed on synthetic cross medium (SCM) plates with and without fertilization (Figure 2a). In the absence of fertilization, the wild-type produced protoperithecia with a relatively uniform diameter (∼100 µm). After fertilization, the wild type produced highly melanized perithecia that were 400 – 600 µm in diameter. In contrast, the unfertilized Δ*set-7* strain produced large and highly melanized protoperithecia structures with sizes ranging from ∼100µm-600µm, overlapping the sizes of both wild-type protoperithecia and fertilized perithecia.

**Figure 2.**
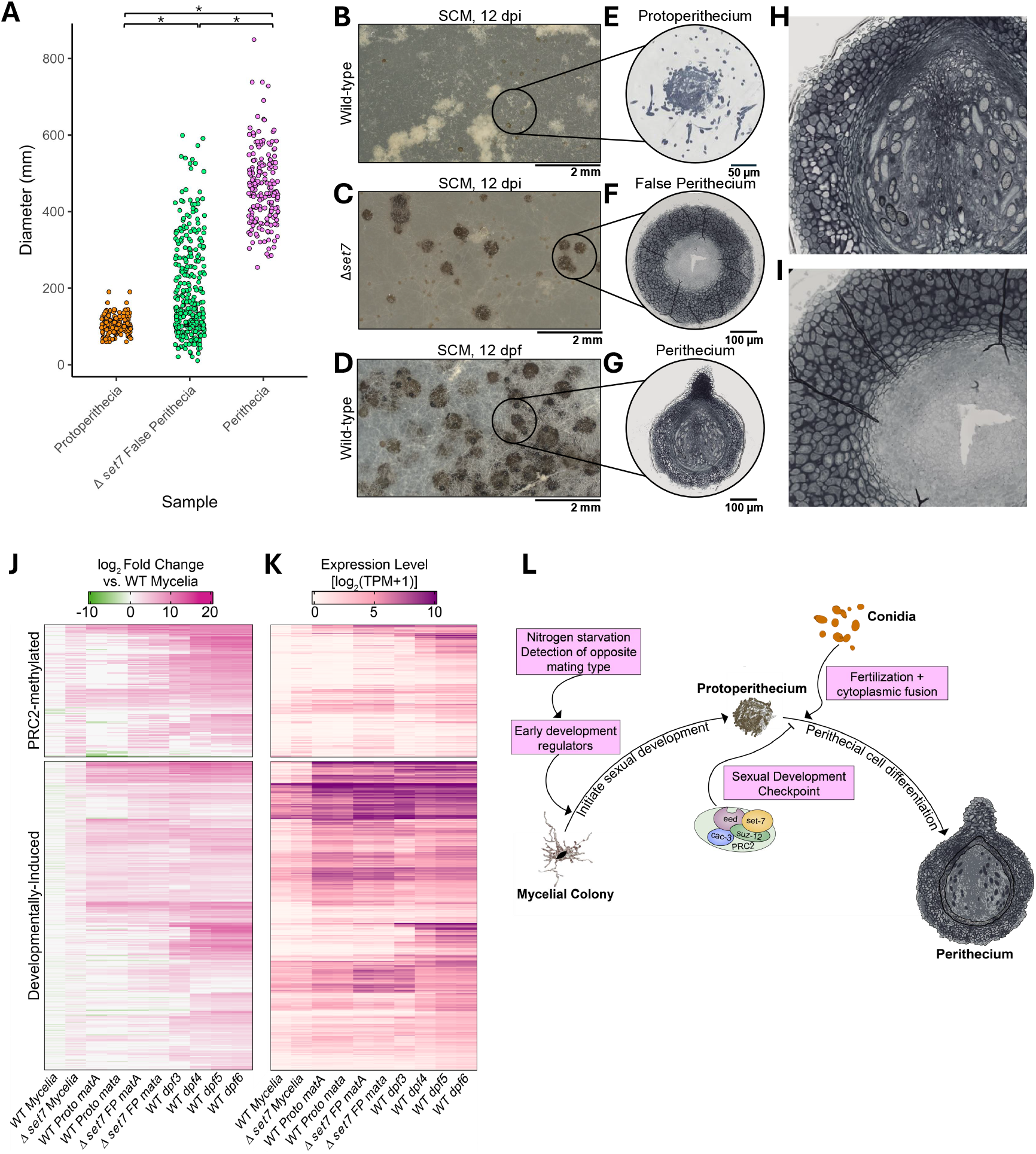
PRC2 is a repressor of perithecial development. (A) The points indicate the diameter (y-axis) of wild-type perithecia, wild-type protoperithecia, or Δ*set-7* false perithecia, as indicated (* = p < 0.001; Kruskal-Wallis test with Holm-adjusted Dunn post-hoc test). (B-D) Representative images are shown for wild-type protoperithecia (unfertilized; 12 days post-inoculation), *Δset-7* false perithecia (unfertilized; 12 days post-inoculation), and wild-type perithecia (12 days post-fertilization) grown on SCM. Scale bars indicate 2 mm. (E-G) Representative images of paraffin-embedded, semi-thin sections of each structure from part A are shown. Sections were stained with toluidine blue before imaging using Bright Field microscopy. The scale bars indicate size in µm. (H-I) The magnified images of perithecia and false perithecia show distinct cell morphologies in the peridium. (J) The heatmaps illustrate relative expression levels (log_2_ fold change relative to mycelial samples) of PRC2-methylated (top) and non-PRC2-methylated DIGs (bottom) for the indicated samples. (K) Raw expression values (TPM+1) are plotted as in J. (L) A model for PRC2-mediated regulation of sexual development is shown. PRC2 represses genes involved in differentiation of perithecial cells until proper signals (fertilization) are received.

To characterize the tissues within the aberrant structures produced by the unfertilized Δ*set-7* strain, we made semi-thin sections through wild-type protoperithecia, wild-type perithecia, and false perithecia from a *Δset-*7 strain and examined the morphology under the microscope (Figure 2b-2i). False perithecia produced by Δ*set-7* differentiated three cell layers: an outer layer of highly melanized tissue, an inner layer of elongated cells, and a layer of small, indistinct cells within the center. The two former cell layers resemble those found in the peridium of a fertilized wild-type perithecium. However, while fertilized perithecia contain ascogenous hyphae in the central cavity, this tissue was absent in false perithecia. Other features of mature perithecia, including the beak and ostiole, were also missing in these structures. We conclude that loss of H3K27me3-mediated repression leads to aberrant development of the peridium, a phenotype resembling homeotic transformations observed in PRC2-deficient plant, insect, and animal cells. A similar transformation was previously observed in the fungus *Podospora anserina*, where loss of H3K27me3 triggers overproduction of male cells (24). Together, these results show that PRC2 is a repressor of perithecial development.

To better understand transcriptional changes underlying this morphological transformation, we extracted RNA from Δ*set-7* strains grown overnight in liquid medium or on solid SCM for 12 days. On SCM plates, wild-type cells produce abundant protoperithecia (PP) and Δ*set-7* strains produce abundant false perithecia (FP). When Δ*set-7* was grown in liquid culture, a subset of H3K27me3-enriched genes exhibited weak upregulation (218/516, 42%) (Figure 2j-k, Figure S2b), consistent with prior studies (7). In contrast, a significant fraction of PRC2-methylated genes (336/516, 65%) were strongly upregulated in FP produced by *Δset-7* (Figure 2j-k, Figure S2b). These genes are typically expressed during perithecial development after fertilization has taken place (Figure 2j-2k). In the FP samples, upregulation of DIGs was not restricted to PRC2-methylated genes, suggesting that global transcriptional reprogramming has occurred in these tissues (Figure 2j-k, Figure S2b). Taken together, our data suggest that PRC2 is required for repression of DIGs that pattern the outer wall of the perithecium.

We propose that PRC2 and its product, H3K27me3, form a developmental checkpoint in *N. crassa* (Figure 2l). Fungi must integrate multiple signals including nutritional stress and fusion with a partner of the opposite mating type to determine if differentiation of fruiting bodies is appropriate. Repressive chromatin assembled by PRC2 functions as a critical checkpoint that ensures appropriate conditions for fruiting body development have been met before terminal differentiation is initiated.

## Data Availability

All RNA-sequencing data generated in this study are freely available at the NCBI Gene Expression Omnibus (https://www.ncbi.nlm.nih.gov/geo/) under the accession GSE296621. The code used for this analysis can be found on GitHub (https://github.com/UGALewisLab/2025_PRC2_Controls_Development).

## Acknowledgements

Sectioning and staining of protoperithecia, false perithecia, and perithecia was performed by Mary Ard at the Georgia Electron Microscopy Core in Athens,GA. We thank Ammal Abduljalil and Adrianna Acosta-Ortiz for contributions to early experiments with false perithecia.

